# Interdependent regulation of alternative splicing by SR and hnRNP proteins

**DOI:** 10.1101/2024.08.19.608666

**Authors:** Megan E. Holmes, Klemens J. Hertel

## Abstract

Alternative pre-mRNA splicing is a combinatorial process involving SR and hnRNP splicing factors. These proteins can silence or enhance splicing based on their expression levels and binding positions. To better understand their combinatorial and interdependent regulation, computational analyses were performed using HepG2 and K562 cell knockdown and binding datasets from the ENCODE Project. Analyses of diMerential splicing for 6 SR proteins and 13 hnRNP knockdowns revealed statistically significant exon overlap among most RBP combinations, albeit at diMerent levels. Neither SR proteins nor hnRNPs showed strong preferences for collaborating with specific RBP classes in mediating exon inclusion. While SRSF1, hnRNPK, and hnRNPC stand out as major influencers of alternative splicing, they do so predominantly independent of other RBPs. Meanwhile, minor influencers of alternative splicing such as hnRNPAB and hnRNPA0 predominantly regulate exon inclusion in concert with other RBPs, indicating that inclusion can be mediated by both single and multiple RBPs. Interestingly, the higher the number of RBPs that regulate the inclusion of an exon, the more variable exon inclusion preferences become. Interdependently regulated exons are more modular and have diMerent physical characteristics such as reduced exon length compared to their independent counterparts. A comparison of RBP interdependence between HepG2 and K562 cells provides the framework that explains cell-type-specific alternative splicing. Our study highlights the importance of the interdependent regulation of alternative exons and identifies characteristics of interdependently regulated exons that diMer from independently regulated exons.

## Introduction

During splicing introns are excised from the pre-mRNA strand and the remaining exons are ligated to form an mRNA strand (Maniatis and Tasic 2002; Woodley and Valcárcel 2002; House and Lynch 2008). Alternative splicing (AS) events are instances where exons are skipped (exon skipping), introns are retained (intron retention), or alternative splice sites are chosen (alternative 3’/5’ selection) (Wagner and Berglund 2014; Woodley and Valcárcel 2002; Kim and Lee 2008). Given the limited number of genes in eukaryotic genomes, AS is essential for diversifying eukaryotic proteomes (Nilsen and Graveley 2010; Manuel et al. 2023). AS is responsible for a large percentage of that diversity, with the majority of human multi-exon genes having AS events (Pan et al. 2008; Wang et al. 2008; Johnson et al. 2003).

Exon skipping (ES) is the most frequent form of AS where changes in exon inclusion generate different mRNA isoforms. As with all forms of AS, ES is a combinatorial process that is regulated by both cis-acting elements and trans-acting factors (Shenasa and Hertel 2019; Douglas and Wood 2011). Two classes of trans-acting factors, the SR protein and hnRNP families, have been shown to affect spliceosomal assembly through similar mechanisms (Maniatis and Tasic 2002; House and Lynch 2008). We hypothesized that it is likely that the SR and hnRNP trans-acting factors interdependently regulate ES. Here, interdependent regulation is defined as regulation through direct and/or indirect effects. Direct effects consider instances where the interaction of an RNA binding protein (RBP) with the pre-mRNA affects exon inclusion. Indirect effects may occur when the altered expression of an RBP influences the expression or splicing pattern of another RBP, which could potentially influence exon inclusion. To investigate the interdependent regulation of ES by these two protein families we turned to the ENCODE project which contains RNA-seq knockdown data and eCLIP binding data for over 300 RBPs in both the human hepatocellular carcinoma cell line, HepG2, and the lymphoblast cell line, K562 (Van Nostrand et al. 2020). Here we utilize the available ENCODE knockdown data from every SR and hnRNP for which data was available in HepG2 and K562 cells to determine the prevalence of interdependent regulation among members of the SR and hnRNP families of splicing regulators.

This multidimensional analysis of interdependence shows that 85% of the 171 possible pairs of evaluated SR proteins and hnRNPs interdependently regulate exon inclusion to a variable degree. Exon inclusion can result from the action of one or several RBPs. Notably, as the number of RBPs governing exon inclusion increases, so does the variability in exon inclusion preferences. Exons regulated interdependently exhibit greater modularity and distinct physical traits, such as shorter length when compared to those regulated independently. These results underscore the significance of interdependent regulation in alternative exon inclusion and pinpoint unique characteristics of such exons that are distinct from independently regulated ones.

## Results

### The influence of individual splicing regulators on exon inclusion

SR and hnRNP splicing regulators have been shown to regulate exon inclusion by modulating the recruitment of spliceosomal components to splice sites (Zhou and Fu 2013; Black 2003; Adams, Rudner, and Rio 1996; Wang and Burge 2008; Graveley and Maniatis 1998). To understand the extent to which RNA binding proteins influence exon recognition, we evaluated how many exon inclusion events were significantly affected by the knockdown of 19 individual splicing regulators in HepG2 cells. For this SR and hnRNP analysis, we took advantage of the public ENCODE dataset which reports on gene expression (Love, Huber, and Anders 2014), alternative splicing differences, and binding for more than 300 RNA binding proteins (Van Nostrand et al. 2016; 2020). The broadness of an SR or hnRNP splicing regulator’s role in exon inclusion varies greatly between the analyzed splicing regulators (Fig. 1A, Supplemental Table 1). For example, 1,192 exons were identified with significant changes in inclusion upon the knockdown of SRSF1 but only 11 exons were identified with changes in inclusion upon the knockdown of hnRNPA2B1.

**Figure 1.**
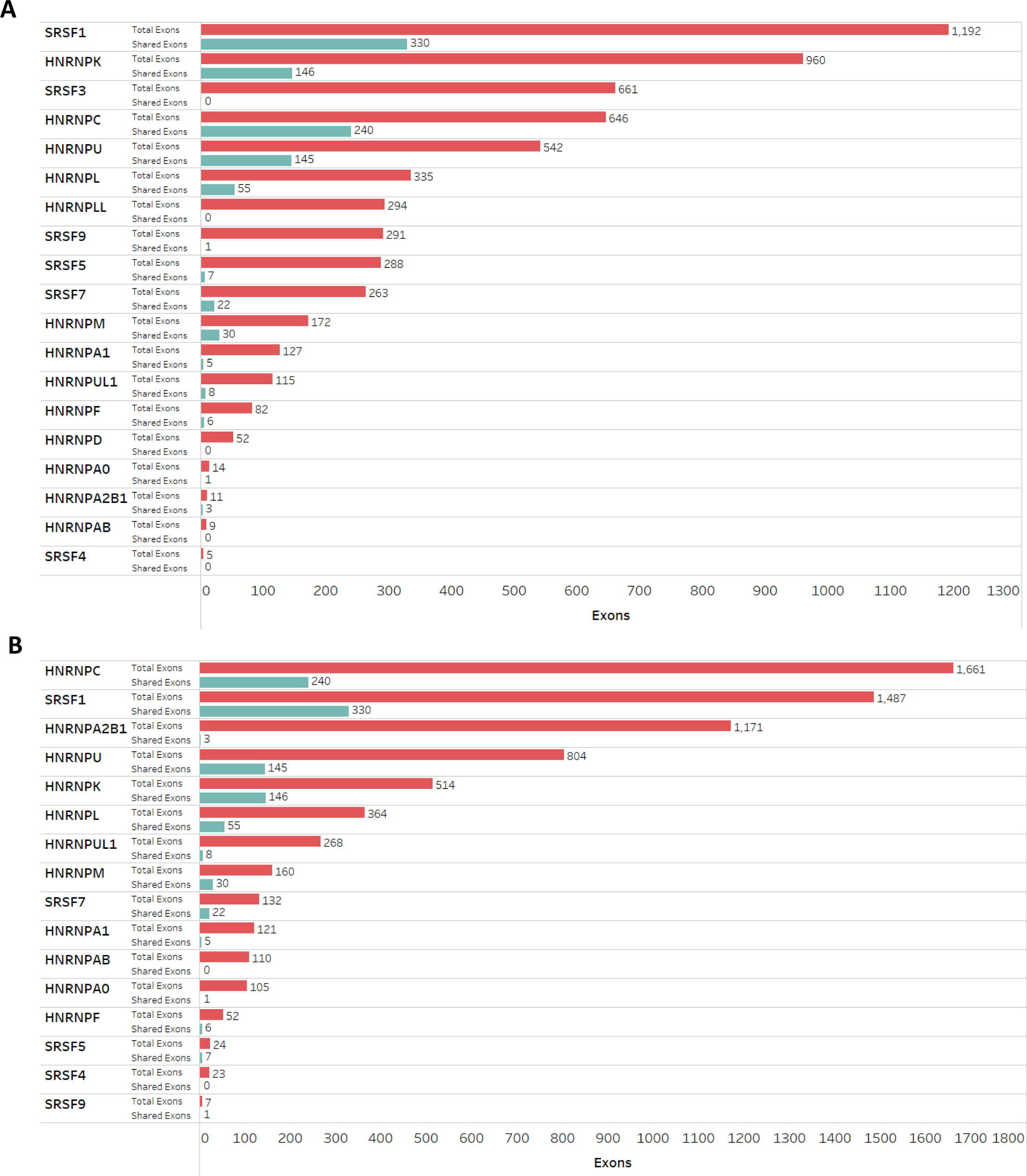
The influence of individual splicing regulators on exon inclusion. A) Bar plot showing the total number of exons (red) with significant changes in exon skipping upon RBP knockdown in HepG2 cells (FDR <= 0.05 and inclusion level difference > 10%). The bar graph is arranged from most (top) to least (bottom) impactful RBP mediating alternative splicing. B) Same as in A), but for K562 cells. Shared exons (teal bars in both panels) are exons with significant changes in inclusion upon RBP knockdown in both HepG2 and K562 cells.

An analogous analysis was performed using all SR and hnRNP knockdown ENCODE datasets available for K562 cells (Fig. 1B, Supplemental Table 2). A comparison between K562 and HepG2 cells demonstrates that the role of an SR or hnRNP splicing regulator can vary greatly between cell lines. While the knockdown of some splicing regulators, such as SRSF1, elicited a comparable number of significant splicing changes in both cell lines, the knockdown of other splicing regulators such as hnRNPA2B1 and hnRNPC displayed drastically different levels of alternative splicing outcomes in HepG2 and K562 cells (Fig. 1B). Interestingly, for each RBP depletion less than 50% of the exons affected were the same between the two cell lines (Fig. 1), demonstrating that the RBPs exhibit cell-type specific differences in splicing targets. We conclude that the extent to which an RBP influences exon inclusion levels varies across the different RBPs and shifts between the two cell lines analyzed.

### SR and hnRNP splicing factors interdependently regulate exon inclusion

Using the alternative splicing information, we evaluated which of the 171 pairs of SR and hnRNP splicing factors available for HepG2 cells exhibit statistically significant interdependent regulation of exon inclusion. Here interdependent regulation is defined as the regulation of an exon by two or more RBPs. For the majority of the RBPs analyzed, interdependently regulated exons make up at least half of the affected exon population indicating a strong likelihood that some of the pairs exhibit statistically significant interdependent regulation (Fig. 2A). The analysis of interdependence between the pairs shows that 85% of the possible splicing regulator pairs exhibit interdependent regulation (Fig. 2B). The amount of interdependent regulation present for exons with increased inclusion and exons with decreased inclusion was similar with 54% and 57% of the possible pairs showing interdependent regulation respectively (Fig. S1). These observations indicate that most SR and hnRNPs interdependently regulate exon inclusion in HepG2 cells. A splicing factor’s level of interdependent splicing regulation varies. This is demonstrated by the difference in the percentage of exons that are also affected by other splicing factors (Fig. 2C). For example, SRSF1-affected exons have low interdependent regulation percentages, while hnRNPA2B1 has higher interdependent regulation percentages (Fig. 2C, rows). However, SRSF1 displays interdependent regulation of splicing with many other splicing regulators while hnRNPA2B1 is more selective for hnRNP proteins in its ability to mediate interdependent exon inclusion (Fig 2C, columns). The data further suggests that the exons affected by hnRNPA2B1 are interdependently regulated by more than one other RBP (Fig. 2D). By contrast, SRSF1 has a larger proportion of exons that are not interdependently regulated, suggesting that SRSF1 modulates exon inclusion mainly independent of other splicing regulators. Interdependent regulation occurs at varying levels between different pairs of splicing regulators without a clear trend of RBP class preferences. This observation is supported by cluster analyses (Fig. S2). RBPs affecting fewer exons appear to exhibit greater levels of interdependent splicing regulation. Qualitatively similar observations were made for RBP interdependence in K562 cells with a slightly lower percentage of pairs exhibiting interdependent regulation (Fig S3A, B) Interestingly, the K652 analysis suggests that most SR proteins exhibit less interdependent regulation in K562 cells when compared to hnRNPs (Fig S3C, D).

**Figure 2.**
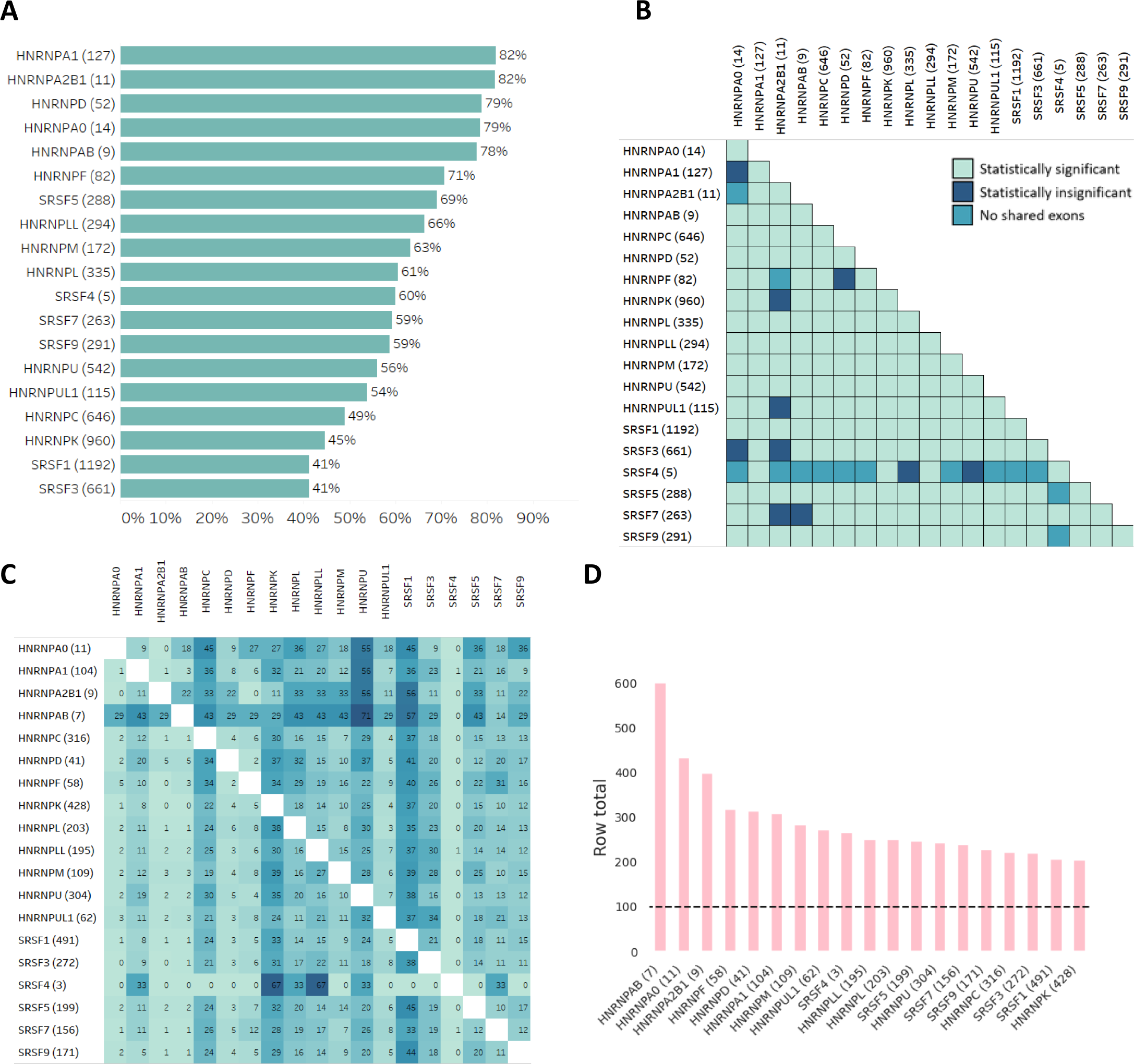
SR and hnRNP splicing factors interdependently regulate exon inclusion. A) Bar plot showing the percentage of each RBP’s total affected exons that are also affected by another RBP. The number in parentheses after each RBP name is the total number of exons with significant changes in exon inclusion upon knockdown of that RBP in HepG2 cells. B) Matrix indicating which of the pairs of the 19 splicing factors analyzed show statistically significant interdependent regulation in HepG2 cells. The number in parentheses after each RBP name is the total number of exons with significant changes in exon inclusion upon knockdown of that RBP in HepG2 cells. Light green indicates statistically significant pairs (p <= 0.05), dark blue indicates statistically insignificant pairs (p > 0.05), and teal indicates no overlap between pairs. c) Heatmap displaying the percentage of alternatively spliced exons that are independently regulated by another RBP in HepG2 cells. The numbers in parentheses are the total number of exons interdependently regulated by that RBP. The values in the cells is the percentage of interdependently regulated exons that are also regulated by the RBP on the horizontal axis. d) Bar plot showing the sum of the values from each row of the heatmap in (c). Values above 100 indicate that some exons are regulated by the RBP of interest and at least one other RBP.

### Enhancing and silencing activity of SR and hnRNPs varies based on interdependence

SR proteins are primarily known to act as splicing enhancers while hnRNP proteins are best known to silence splicing (Graveley 2000; Tange et al. 2001; Maniatis and Tasic 2002; House and Lynch 2008). However, it is also appreciated that splicing regulators can act as both splicing silencers and enhancers, presumably dictated by their binding relative to regulated splice sites (Zahler et al. 2004; Graveley, Hertel, and Maniatis 1999; Krecic and Swanson 1999; Ule et al. 2006; Carranza, Shenasa, and Hertel 2022; Erkelenz et al. 2013; Van Nostrand et al. 2020). To determine whether SR and hnRNP proteins preferentially act as silencers or enhancers, we computed the percentage of exons displaying an increase or decrease in inclusion level upon RBP knockdown (Fig. 3). This analysis was carried out for three different populations of alternatively spliced exons: the total population consisting of every exon affected by the knockdown of the RBP of interest, the interdependent population consisting of all exons whose inclusion is affected by the RBP of interest and at least one other RBP, and the independent population consisting of all exons that are only affected by the RBP of interest. In agreement with previous results, all splicing regulators exhibit both silencing and enhancing activity on the total population albeit with different preferences (Ule et al. 2006; Erkelenz et al. 2013). For example, knockdowns of most SR splicing regulators resulted in preferential exon inclusion loss, supporting the notion that SR proteins primarily act as splicing enhancers (Fig. 3A). The inverse is observed for the majority of hnRNP splicing regulators, consistent with their general role as splicing silencers (Fig. 3B). Remarkably, the tendencies of all SR proteins analyzed shifted from preferential enhancing to preferential silencing when exon inclusion is mediated by at least one other RBP (interdependent population). The splicing-enhancing function of SR proteins is amplified if it is the only RBP acting on exon inclusion (independent population) (Fig. 3A).

**Figure 3.**
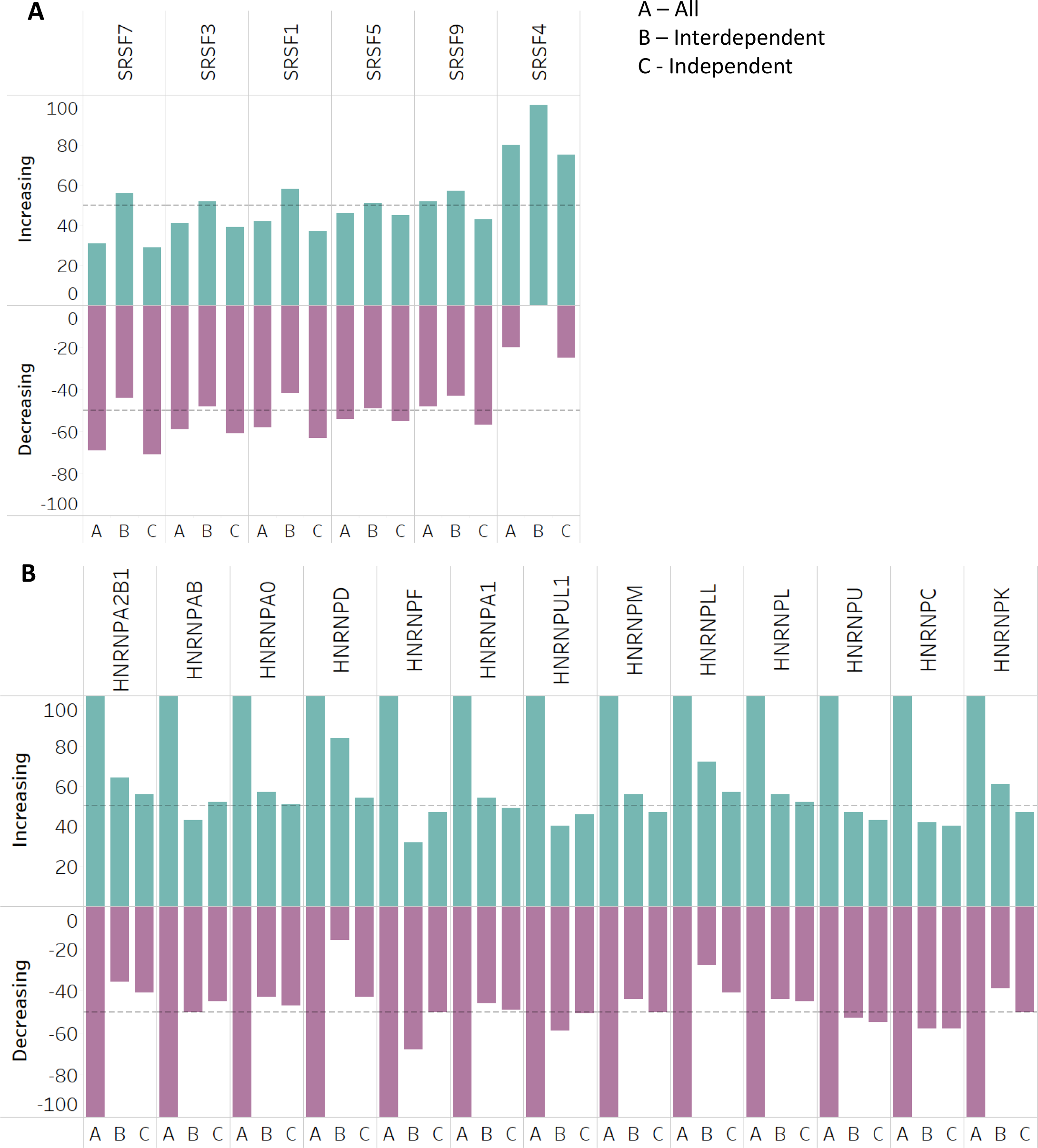
Enhancing and silencing activity of SR and hnRNP proteins varies based on interdependence. A) Grouped bar graph showing the percentage of exons with increasing (positive y-axis) and decreasing (negative y-axis) inclusion levels upon knockdown of individual SR proteins in HepG2 cells. For each RBP three groups are considered. (All) includes every exon with a biologically significant change in inclusion level upon knockdown of the RBP, (Inter) refers to all interdependently regulated exons, and (Ind) includes all independently regulated exons. The dashed lines indicate the 50% mark of increasing (positive) and decreasing (negative) exon populations. B) The same as (A) but for hnRNP proteins in HepG2 cells.

Most of the hnRNPs analyzed exhibited preferential silencing activity, especially within the independent exon population. With minor deviations, these enhancing and silencing preferences are also observed for the RBP knockdown analysis in K562 cells (Fig. S4). We conclude that SR and hnRNP splicing regulators differentially affect exon inclusion levels, but their splicing/enhancer tendencies shift depending on whether the exon is interdependently regulated or not.

### Independently and interdependently regulated exon populations have different characteristics

Typically, exons are highly or lowly included with few having intermediate inclusion levels (∼50%) (Shepard et al. 2011). Considering the difference in the enhancing/silencing activity of the interdependent and independent RBP populations (Fig. 3) we evaluated whether the wildtype exon inclusion levels of RBP-affected exons differ between the two populations. Interestingly, while the average inclusion levels of both groups were similar, the distribution of each group was observed to be considerably different. The independent population exhibited the expected bimodal distribution of lowly and highly included exons (Shepard et al. 2011) while the interdependent group exhibited a unimodal distribution (Fig. 4A, B).

**Figure 4.**
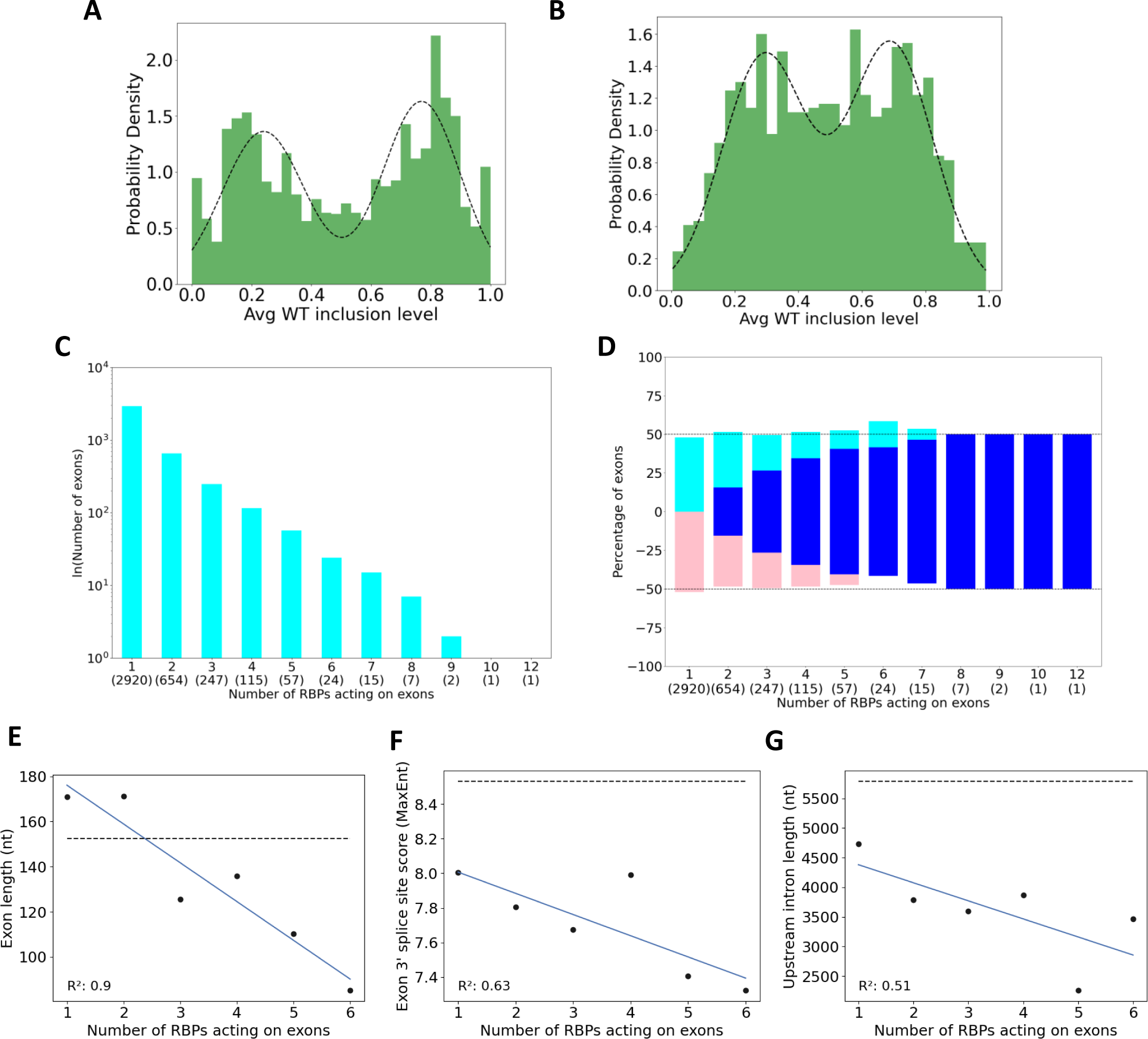
Independently and interdependently regulated exon populations have different characteristics. A) Histogram displaying the distribution of wildtype inclusion level of every independently regulated exon with biologically significant changes in exon inclusion upon knockdown of any RBP in HepG2 cells. B) Histogram displaying the distribution of wildtype inclusion level of every interdependently regulated exon with biologically significant changes in exon inclusion upon knockdown of any RBP in HepG2 cells. C) Bar graph showing the number of exons that are affected by the knockdown of one or multiple RBPs in HepG2 cells. D) Stacked histogram showing how the number of RBPs affecting individual exons dictates preferential increase (teal), decrease (pink), or variable (dark blue) exon inclusion. E) Correlation between exon length and the number of RBPs affecting individual exons. The dashed line indicates the average exon length for all internal exons in the human transcriptome. F) The same as (E) but for the average exon 3’ splice site score. G) The same as (E) but for the average upstream intron length.

To further break down the differences between the independent and interdependent populations, we determined how many RBPs act on an exon by combining the exon inclusion data for all knockdown datasets and computing how many RBPs can act on each exon. Greater than 70% of alternatively spliced exons are only affected by one RBP (independently regulated). Astonishingly, up to 12 RBPs can affect the inclusion level of a single exon showing that some exons experience interdependent regulation with multiple RBPs (Fig. 4C). Within these populations we determined the percentage of exons with an exclusive increase or decrease in inclusion level, or the percentage of exons with inclusion levels in either direction (Fig. 4D). The data shows no preference for enhancing or silencing exon inclusion across the populations. This is expected considering that the effects of hnRNPs and SR proteins are combined in this analysis. As the number of RBPs affecting an exon’s inclusion level increases, the fraction of the population that mediates both increased and decreased exon inclusion becomes increasingly dominant (Fig. 4D). Thus, the cooperation of multiple RBPs in mediating exon inclusion reduces the preferential enhancement or repression of exon inclusion that is typically induced by the actions of a single RBP. Furthermore, the proportion of silenced exons slightly increases as the number of RBPs affecting the inclusion of an exon increases. Similar observations were made from the K562 analysis (Fig. S5A, B).

To determine whether exons that are regulated by a different number of RBPs display unique regulatory features, we compared exon length, intron length, splice site scores, and sequence conservation across the various exon populations. The most significant dependency we observed was a negative correlation between the number of RBPs affecting exon inclusion and exon length (Fig. 4E).

Longer exons are preferentially regulated by a single RBP and as the number of RBPs modulating exon inclusion increases, exon size decreases. Weaker negative correlations were also found between the number of RBPs affecting exon inclusion and the 3’ splice site score strengths as well as the upstream intron length (Fig. 4F, G). In general, splice site scores are lower than the genome average across all populations suggesting that the affected exons are weak. Interestingly, the inclusion of exons with weaker splice sites requires a higher number of interdependent RBPs (Fig. 4F). Compared to the genome average the sequence conservation of all populations is elevated, trending towards increased conservation as the number of RBPs affecting exon inclusion increases. These observations suggest that exons targeted by multiple splicing regulators have evolved to establish a sub-optimal exon recognition potential. Other features such as 3’ splice site score, downstream intron length, and PhyloP score showed insignificant levels of linear correlation (Fig. S5). The analogous K562 analysis yielded similar results (Fig. S6).

## Discussion

In this study, interdependence was defined by the overlap of exons between the differential splicing analysis of two RBP knockdowns. The hypothesis that SR/hnRNPs interdependently regulate exon inclusion is supported by this study as 85% of the 171 possible non-duplicate pairs of the SR/hnRNPs available for HepG2 cells in the ENCODE dataset exhibit statistically significant interdependent regulation. A similar extent of interdependent regulation was observed in K562 cells indicating that while there are variations in the other aspects analyzed, interdependent regulation is not unique to HepG2 cells. The broad range in the number of exons that an RBP regulates and the variation of those counts per RBP between cell lines suggests that each cell line has its own set of more prevalent regulators. SRSF1 and hnRNPC emerged as two of the most influential splicing regulators as their knockdown resulted in differential exon inclusion for many exons in both cell lines (Fig. 1). The exons affected by lowly active RBPs have higher levels of interdependent regulation suggesting that lowly active RBPs may only function in those cell lines as interdependent regulators. A surprising observation was the fact that >70% of exons undergoing alternative splicing upon RBP knockdown appear independently regulated, meaning their inclusion level was only affected by the knockdown of one SR or hnRNP protein (Fig. 4C). While other splicing regulators not evaluated in this study could affect such independently regulated exons, a picture emerges that many alternatively spliced exons are predominantly regulated by the activity of one splicing regulator (Fig. 5). A case can be made that the involvement of multiple RBPs in regulating alternative exon inclusion allows the fine-tuning of exon inclusion levels and increased modularity in mediating increased or decreased exon inclusion.

**Figure 5.**
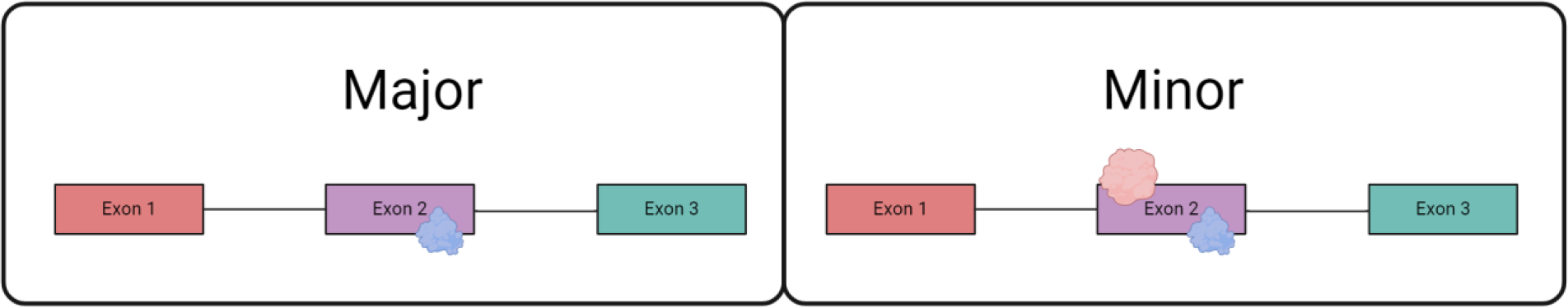
Mechanisms of RBP-regulated exon inclusion. Models for regulatory mechanisms of exon inclusion. The majority of alternatively spliced exons are independently regulated, meaning that only one RBP primarily influences exon inclusion. At a lower frequency, multiple RBPs influence exon inclusion through interdependent regulation. Interdependent regulation can be a cooperation between two or many different RBPs.

### Pairs of RBPs, family of RBPs, and trends in interdependent regulation

The interdependence matrix (Fig. 2C) illustrates what frequency exons affected by the knockdown of an RBP defined on the Y-axis are also affected by the knockdown of RBPs defined by the X-axis. Crucial in evaluating the information presented is considering the total number of interdependently regulated exons that are associated with the knockdown of an RBP. For example, the knockdown of hnRNPA1 changes the inclusion for 127 exons (Fig. 1A), 104 of which (82%) (Fig. 2C) are also regulated by other RBPs. The matrix shows that 56% of hnRNPC’s interdependently regulated exons (58 exons) are also regulated by hnRNPU. Thus, hnRNPC and hnRNPU frequently co-regulate exon inclusion. On the other hand, only 6 of hnRNPL’s 203 affected exons (3%) are interdependently affected by hnRNPUL1, suggesting that hnRNPL and hnRNPUL1 do not operate frequently on the same set of exons. The matrix also demonstrates that the degree of interdependent regulation between RBPs is not defined by family traits nor is it influenced by whether an RPB is a preferential enhancer or repressor of exon inclusion (Fig. S2). Thus, the SR and hnRNP splicing regulators display a flexible potential to contribute to exon inclusion.

### Direct versus indirect effects mediating interdependent splicing regulation

Splicing regulators are known to bind the pre-mRNA and mediate splicing activity by influencing the recruitment efficiency of spliceosomal components. Thus, when alternative splicing is detected upon knockdown of a splicing regulator, it is assumed that the difference in an exon’s inclusion level is caused by the loss of the direct interaction between the pre-mRNA and the slicing regulator. While this “direct effect” is logical, it is also possible that indirect effects, such as induced changes in the expression of other splicing regulators could trigger alternative splicing. The available ENCODE data (splicing, transcriptomics, eCLIP binding) provided only limited insights into addressing to what degree the observed splicing changes upon RBP knockdown were caused by direct (loss of RBP binding) or potential indirect effects (altered expression or splicing of other RBPs). Only a small overlap between the stringently filtered high-confidence eCLIP peak dataset from the ENCODE project and alternatively spliced exons was observed (Fig. S7) (Van Nostrand et al. 2020), providing only limited circumstantial evidence in support of direct effects. Even if the overlap was significantly greater, a correlation between the presence of eCLIP binding peak within and around alternatively spliced exons does not prove an RBP’s involvement in regulating exon inclusion as the RBPs may be bound to exons before or after splicing occurs. Furthermore, the absence of eCLIP binding does not disprove direct binding, especially when RBP binding occurs within short-lived introns. Nevertheless, direct effects are still highly likely to mediate observed alternative splicing, particularly in the case of exons affected only by a single RBP (independently regulated exons). Transcriptomic and alternative splicing analyses demonstrated that the knockdown of an RBP can cause changes in the expression or splicing pattern of the other RBPs, raising the potential that indirect effects impact alternative splicing outcomes of RBP knockdowns (Fig. S8). While it is unknown to what degree these changes in other RBP expression and isoform representation affect alternative splicing, they must be considered as potentially influential when evaluating the alternative splicing landscape of single RBP knockdowns. Thus, alternative splicing changes observed upon RBP knockdown are caused by both direct and indirect effects.

### Features of interdependently regulated exons

The bimodal distribution of exon inclusion levels for the independently regulated exons is similar to the exon inclusion distribution of human exons, with an underrepresentation of intermediate exon inclusion levels (Shepard et al. 2011). Interestingly, the exon inclusion distribution of interdependently regulated exons is strikingly different as intermediate inclusion level exons are overrepresented (Fig. 4A, B). These exons are also more easily shifted towards inclusion or exclusion than independently regulated exons. This increased modularity likely makes it easier for the balance of the exon’s inclusion to be easily shifted in either direction by the combined regulation of RBPs. Contrary to expectations based on steric hindrance, shorter exons are associated with an increased number of RBPs regulating its inclusion (Fig. 4E), perhaps through a combination of exonic and intronic binding sites.

### Cell type specificity

From the ENCODE project we were able to analyze interdependent regulation in both HepG2 and K562 cells (Van Nostrand et al. 2020). While many general outcomes were qualitatively similar, our analysis showed extensive cell type specificity. For example, the degree of regulation mediated by hnRNPA2B1 varied significantly between the two cell lines (Fig. 1) even though the expression difference between the two cell lines was greater than 2-fold (Fig. S9). Along with differences in RBP activity between the cell lines, the regulated exons had minimal overlap across the two cell lines (Fig. 1). These observations suggest that different gene expression programs in HepG2 and K562 cells result in RBP activity and exon selection. Alternatively, the abundance of RBPs in each cell line could alter exon selection. While modest, differential gene expression of the RBPs evaluated in each cell line supports that latter argument (Fig S9). Thus, while an RBP’s ability to interdependently regulate splicing remains constant, cell-type specific differences in alternative splicing appear to be mediated through altered expression of splicing regulators and target genes.

## Materials and Methods

### Identifying evidence of indirect versus direct effects of interdependence

Our analysis of interdependent regulation was carried out on 19 pre-mRNA splicing factors consisting of 6 SR proteins and 13 hnRNPs using HepG2 data from the ENCODE dataset (Van Nostrand et al. 2020). An analogous analysis was done using the K562 data for all 17 available SR/hnRNPs. We considered two metrics of knocking down an RBP as potential evidence for indirect effects. First, if the knockdown of an RBP induces changes in the expression of other RBPs, and second if the knockdown of an RBP induces changes in the splicing pattern of other RBPs. Des-seq was used to address the first potential inducer of indirect effects using ENCODE datasets with a p-value <= 0.05 cutoff. rMATS was used to determine whether the knockdown of an RBP resulted in differential splicing of other SR/hnRNPs. Significant splicing pattern changes in SR/hnRNPs resulting from the knockdown of another SR/hnRNP were characterized as instances where an RBP had at least one exon that exhibited statistically significant (FDR <= 0.05) change and a > 10% difference in inclusion.

### Identifying RBP pairs that exhibit statistically significant interdependent regulation

To identify statistically significant interdependent regulation between each of the 171 possible non-duplicate RBP pairs we used the survival function:

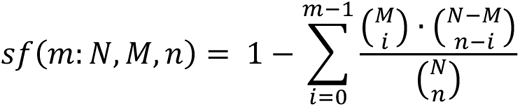

- m: the number of exons present in both filtered rMATS files for a given pair.
- N: the total number of exons in the unfiltered rMATS files combined for the proteins of interest
- M: the total number of exons in the filtered rMATS file for the first RBP of interest
- n: the total number of exons in the filtered rMATS file for the second RBP of interest

Exon skipping rMATS files comparing wildtype HepG2 and the HepG2 SR/hnRNP knockdowns were filtered for exons with an FDR <= 0.05 and an inclusion level difference > 10% to create the ‘filtered’ rMATS files. ‘Unfiltered’ rMATS files refer to the raw files without duplicates. Pairs whose survival function returned a p-value <= 0.05 exhibit statistically significant interdependent regulation.

### Clustering of RBP interdependence activity

Clustering of the interdependence matrix (Fig. 2C) was carried out using Clustergrammer (Fernandez et al. 2017). The rows were hierarchically clustered, using the Scipy library in Python, using Jaccard distance and average linkage. The column order was ranked by column sum. The intensity of the red cells in the output heatmap represents the magnitude of values in the input matrix (Fig. S2).

### Identifying evidence of direct effects

To identify evidence for potential direct effects between RBP binding and exon inclusion changes, we cross-referenced eCLIP direct binding data with alternative splicing outcomes. eCLIP data from the ENCODE dataset for the 10 analyzed RBPs for which eCLIP data was available (hnRNPA1, hnRNPC, hnRNPK, hnRNPL, hnRNPM, hnRNPU, hnRNPUL1, SRSF1, SRSF7, and SRSF9) was used to identify overlaps with rMATS-generated alternative splicing information. An overlap was defined if eCLIP signals of RBP pairs were identified within alternatively spliced exons and 150bp of upstream and downstream intronic sequence. The stringent eCLIP peak detection metric as defined by the ENCODE consortium was used (Van Nostrand et al. 2020). For each possible non-duplicate pair for these 10 RBPs, we used PyRanges to find any instances in which there were eCLIP peaks within any of the exons or their 150bp upstream and downstream intronic regions (Stovner and Sætrom 2020). Exons analyzed were present in both knockdown files of the RBPs in the pair and were filtered for FDR <= 0.05 and > 10% inclusion level difference.

### Correlating exon features with exons affected by varying numbers of RBP knockdowns

Alternatively spliced exons were binned by the number of RBPs affecting their inclusion. Exon parameters such as exon length, the upstream and downstream intron lengths, the 3’ and 5’ splice site strengths, and sequence conservation (PhyloP score) were extracted from UCSC genome browser tracks and from MaxEnt splice site score calculators (Eng et al. 2004). The averages of these characteristics were then compared across each exon bin. A linear regression model was used to determine if any of the characteristics exhibited a linear correlation across groups with increasing numbers of affecting RBPs.

## Acknowledgments

We would like to thank the Hertel lab for helpful conversations. This work was supported by grants from the NIH (R35 GM145254 and R01 GM062287 to K.J.H)

